# Nomenclature updates to the hemagglutinin gene clade designations resulting from the continued evolution of high pathogenicity avian influenza A(H5) virus clades 2.3.2.1c and 2.3.4.4

**DOI:** 10.1101/2025.11.23.690055

**Authors:** Tommy T. Lam, C. Todd Davis, World Health Organization/World Organisation for Animal Health/Food and Agriculture Organization (WHO/WOAH/FAO) H5 Evolution Working Group

## Abstract

The evolutionary divergence of the A(H5) hemagglutinin (HA) gene of high pathogenicity avian influenza (HPAI) viruses (A/goose/Guangdong/96 lineage) was analyzed by phylogenetic and average pairwise distance methods to identify clades that merit nomenclature updates. Based on this assessment, 12 new clade designations were recommended based on division of clade 2.3.2.1c and 2.3.4.4 viruses, which were reported in Africa, Antarctica, Asia, Europe, the Middle East, the Americas and Oceania since the most recent WHO/WOAH/FAO H5 Evolution Working Group update.

## INTRODUCTION

The hemagglutinin (HA) gene of high pathogenicity avian influenza (HPAI) A(H5) viruses derived from the A/goose/Guangdong/1/96 (gs/GD/96) A(H5) HA lineage has continued to evolve since previous updates to the A(H5) clade nomenclature by the WHO/WOAH/FAO H5 Evolution Working Group in 2014 (1–5). Since the start of 2015, which corresponded to the date of the last update, HPAI A(H5) viruses have continued to circulate in both poultry and wild birds across regions of Africa, Asia, Europe, and the Middle East and subsequently spread to the Americas and Antarctica (6,7). Clade 2.3.2.1a viruses have consistently been detected in poultry in Bangladesh, India and Nepal, and are associated with human infections in Australia (travel-associated case from India), Bangladesh, India and Nepal (8). Clade 2.3.2.1c viruses were detected in avian hosts in numerous countries including Bulgaria, China, Indonesia, India, Japan, Lebanese Republic, Malaysia, Myanmar, Romania, Russian Federation, and several West African countries between 2015 to 2018, and have circulated persistently in the Greater Mekong subregion (Cambodia, Lao People’s Democratic Republic, Vietnam) since 2014. While other clades of A(H5) have been displaced over time, clade 2.3.2.1c has remained entrenched in poultry in the Asian region but recent reassortment occurring with clade 2.3.4.4 viruses has resulted in novel genotypes detected in the region (9). While persistently circulating viruses belonging to clades 2.3.2.1a and 2.3.2.1c have retained N1 NA genes, clade 2.3.4.4 viruses became the globally predominant clade and carried numerous NA genes, including N1, N2, N3, N4, N5, N6, and N8 (8). Since 2015, in addition to the extensive intercontinental spread of clade 2.3.4.4 A(H5N1) viruses, A(H5N2), A(H5N6) and A(H5N8) subtype viruses belonging to clade 2.3.4.4 have also spread globally (8, 10).

This extensive geographic spread underscores the dynamic evolution of clade 2.3.2.1c and 2.3.4.4 A(H5) viruses, with recent studies highlighting their genetic divergence within and across temporal and geographic regions, as well as among viruses circulating in wild birds and poultry and those that have spilled over to mammalian species in many regions including dairy cattle in the USA. In addition, the emergence of new genetic groups has coincided with continued reports of human infections. Clade 2.3.2.1c A(H5N1) viruses have been linked to human cases in China, Nepal, Cambodia, Indonesia, and Vietnam, while clade 2.3.4.4 viruses, including A(H5N1) (e.g., in the United States) and A(H5N6) (e.g., in China), have also caused human infections (8).

This report describes the phylogenetic analysis of gs/GD/96 lineage A(H5) HA sequence data available since the last nomenclature update at the end of 2014 (new data for this report was deposited in databases from 1 January 2015 through 04 July 2024) and nomenclature designations for the emerged clades proposed by the WHO/WOAH/FAO H5 Evolution Working Group.

## MATERIALS and METHODS

### Sequence alignments

A total of 18,191 A(H5) HA sequences from GISAID and GenBank databases with virus sequence deposit dates up to and including 04 July 2024 were used in this nomenclature analysis. Sequences were curated using custom Perl scripts and database filters (available upon request) as in the previous nomenclature updates whereby sequences were removed before further analysis if analysis detected signs of recombination (11), duplicates, more than 5 ambiguous nucleotides, less than 60% alignment length, or frameshifts. Data were aligned via MAFFT v7.388 (12) and trimmed to the beginning of the mature A(H5) HA protein gene sequence using BioEdit 7.2.5.

### Phylogenetic trees

Approximate maximum likelihood trees (GTR+GAMMA with 10,000 resamples for local support of topology) were constructed using FastTree v2.1.11 (13). Automated clade annotation of new sequences used the LABEL H5v2015 classification module. Pairwise p-distance matrices were calculated in MEGA5.1 (15) and group averages were calculated with a custom Perl script. Figure 1 is a tree of 261 representative A(H5) HA genes rooted to gs/GD/96. All sequence names, their assigned clades, accession numbers, and data sources are provided in Supplementary Data S2-3.

**Figure 1.**
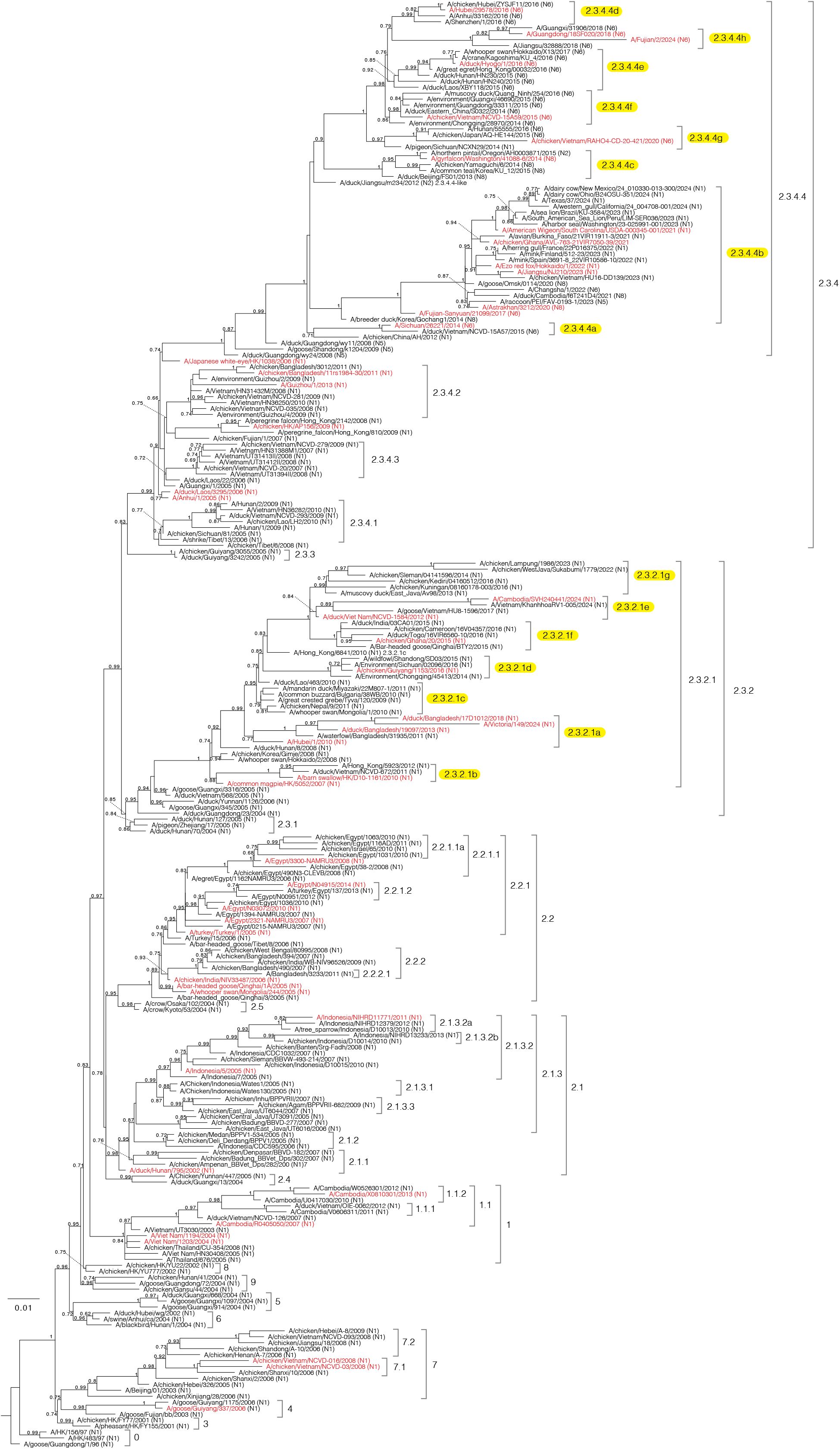
Phylogenetic relationships of recently diverged A/goose/Guangdong /1/1996 (Gs/GD/96)-like H5 hemagglutinin (HA) genes. A maximum likelihood tree of 242 HA nucleotide sequences from Gs/GD/96 lineage of HPAI H5 viruses was constructed with 10,000 resamples for local support of topology (above branches) using FastTree2 (GTR+GAMMA) and rooted to Gs/GD/96. Selected sequences are representative of all clades to render a condensed but accurate phylogenetic topology while including viruses from diverse countries, vaccine candidates (strain names highlighted in red), and human cases. All viruses are A(H5N1) unless labeled otherwise. Newly designated clades are highlighted in yellow. Scale bar denotes nucleotide substitutions per site. Sequence accession numbers are provided in Supplementary Data S2.

Clade designations for newly emerged phylogenetic groups were recommended according to previously established criteria (4). Briefly, new clade designations required presence of viruses sampled after 2015, formation of monophyletic clusters with local support values ≥60%, and within-clade average pairwise distances of ≤1.5%; however this last upper limit is relaxed when phylogenetic and surveillance data are insufficient to support the split of the clade (1). New clades were evaluated in the context of two major phylogenetic groups identified: 2.3.2.1c and 2.3.4.4. As used for the previous update to the A(H5N1) nomenclature system (1), fifth-order groups are designated using an additional letter to the right of the fourth-order clade (i.e., 2.3.2.1c).

## RESULTS

The reconstructed phylogenies revealed monophyletic groups in the majority of circulating clades. Existing clades containing new sequences were analyzed to determine the within-clade average pairwise nucleotide distances (APD) (Figure 1 and Supplementary Figure 1A,B). Based on previously defined nomenclature criteria, we concluded that clades 2.3.2.1c and 2.3.4.4 required splitting (Table 1). Clade 2.3.2.1c had an internal APD of 3.93% with viruses circulating in Indonesia, India, West Africa, China, Vietnam since 2015. This clade was subsequently divided into new subclades 2.3.2.1d through 2.3.2.1g, with an average intra-clade APD of 2.04%. Viruses circulating in China and Vietnam from 2012–2017 remained classified under clade 2.3.2.1c, while those diverging between 2014–2016 formed a separate lineage designated as clade 2.3.2.1d. A group of viruses identified in Vietnam and later in Cambodia during 2020–2024 was classified as clade 2.3.2.1e. Viruses spreading to India, the Middle East, and Africa between 2013 and 2017 were categorized under clade 2.3.2.1f. Meanwhile, viruses circulating in Indonesia from 2012–2016, previously grouped with other Asian sequences under clade 2.3.2.1c, were reassigned to a distinct subclade, 2.3.2.1g. This classification reflects the monophyletic nature of the Indonesian sequences, suggesting localized circulation during that period.

**Table 1.** Genetic divergence within WHO/WOAH/FAO H5 clades since January 1, 2015.

| Current clade | # of sequences | intra-clade APD | nearest clade for comparison | inter-clade APD | New clade designation | # of sequences | intra-clade APD | nearest clade for comparison | inter-clade APD |
| --- | --- | --- | --- | --- | --- | --- | --- | --- | --- |
| 0 | 7 | 1.82% | 5 | 4.89% | 2007* |  |  |  |  |
| 1 | 33 | 1.29% | 9 | 3.40% | 2010 |  |  |  |  |
| 1.1 | 10 | 1.25% | 1 | 2.54% | 2008 |  |  |  |  |
| 1.1.1 | 7 | 1.77% | 1.1 | 3.12% | 2012 |  |  |  |  |
| 1.1.2 | 15 | 1.66% | 1.1 | 3.81% | 2014 |  |  |  |  |
| 2.1.1 | 15 | 1.97% | 2.5 | 2.87% | 2007 |  |  |  |  |
| 2.1.2 | 26 | 1.09% | 2.1.1 | 2.69% | 2006 |  |  |  |  |
| 2.1.3 | 14 | 1.65% | 2.1.1 | 3.27% | 2007 |  |  |  |  |
| 2.1.3.1 | 14 | 1.15% | 2.1.1 | 3.22% | 2011 |  |  |  |  |
| 2.1.3.2 | 64 | 1.84% | 2.1.1 | 3.71% | 2010 |  |  |  |  |
| 2.1.3.2a | 63 | 2.16% | 2.1.3.2b | 4.46% | current Indonesian virus |  |  |  |  |
| 2.1.3.2b | 52 | 1.95% | 2.1.3.2a | 4.46% | 2016 |  |  |  |  |
| 2.1.3.3 | 22 | 1.31% | 2.1.1 | 3.42% | 2010 |  |  |  |  |
| 2.2 | 21 | 1.16% | 2.2.1 | 1.86% | 2008 |  |  |  |  |
| 2.2.1 | 15 | 1.35% | 2.2 | 1.85% | 2013 |  |  |  |  |
| 2.2.1.1 | 8 | 1.75% | 2.2.1.1a | 2.46% | 2010 |  |  |  |  |
| 2.2.1.1a | 8 | 1.70% | 2.2.1.1 | 2.46% | 2011 |  |  |  |  |
| 2.2.1.2 | 70 | 1.78% | 2.2.1 | 2.95% | current Egypt virus |  |  |  |  |
| 2.2.2 | 9 | 1.27% | 2.2 | 2.31% | 2010 |  |  |  |  |
| 2.2.2.1 | 8 | 1.36% | 2.2.2 | 1.97% | 2011 |  |  |  |  |
| 2.3.2.1 | 4 | 1.22% | 2.3.2.1b | 3.41% | 2011 |  |  |  |  |
| 2.3.2.1a | 385 | 2.85% | 2.3.2.1 | 5.73% | current Bangladesh virus |  |  |  |  |
| 2.3.2.1b | 20 | 1.42% | 2.3.2.1 | 3.43% | 2013 |  |  |  |  |
| 2.3.2.1c | 1062 | 3.93% | 2.3.2.1 | 4.89% | 2.3.2.1c | 177 | 1.23% | 2.3.2.1e | 4.32% |
|  |  |  |  |  | 2.3.2.1d | 53 | 1.02% | 2.3.2.1c | 3.33% |
|  |  |  |  |  | 2.3.2.1e | 450 | 3.31% | 2.3.2.1f | 3.92% |
|  |  |  |  |  | 2.3.2.1f | 157 | 1.70% | 2.3.2.1e | 3.92% |
|  |  |  |  |  | 2.3.2.1g | 225 | 3.16% | 2.3.2.1e | 4.41% |
| 2.3.4 | 19 | 2.63% | 2.3.4.3 | 2.69% | 2010 |  |  |  |  |
| 2.3.4.1 | 16 | 1.12% | 2.3.4.3 | 3.79% | 2010 |  |  |  |  |
| 2.3.4.2 | 9 | 1.80% | 2.3.4.3 | 2.85% | 2013 |  |  |  |  |
| 2.3.4.3 | 34 | 0.92% | 2.3.4 | 2.69% | 2009 |  |  |  |  |
| 2.3.4.4 | 15975 | 3.50% | 2.3.4 | 9.96% | 2.3.4.4 | 12 | 1.70% | 2.3.4.4-like | 4.91% |
|  |  |  |  |  | 2.3.4.4a | 113 | 1.09% | 2.3.4.4-like | 5.29% |
|  |  |  |  |  | 2.3.4.4b | 13877 | 1.98% | 2.3.4.4-like | 7.36% |
|  |  |  |  |  | 2.3.4.4c | 862 | 1.68% | 2.3.4.4-like | 5.07% |
|  |  |  |  |  | 2.3.4.4d | 97 | 0.87% | 2.3.4.4f | 2.61% |
|  |  |  |  |  | 2.3.4.4e | 295 | 1.54% | 2.3.4.4-like | 2.52% |
|  |  |  |  |  | 2.3.4.4f | 93 | 1.35% | 2.3.4.4-like | 2.31% |
|  |  |  |  |  | 2.3.4.4g | 137 | 2.59% | 2.3.4.4-like | 3.73% |
|  |  |  |  |  | 2.3.4.4h | 489 | 2.67% | 2.3.4.4-like | 4.44% |
| 2.5 | 2 | 0.19% | 2.2 | 2.40% | 2006 |  |  |  |  |
| 5 | 3 | 0.84% | 9 | 4.21% | 2009 |  |  |  |  |
| 7 | 10 | 3.58% | 5 | 5.93% | 2008 |  |  |  |  |
| 7.1 | 3 | 2.81% | 7 | 4.57% | 2008 |  |  |  |  |
| 7.2 | 4 | 2.69% | 7 | 5.67% | 2014 |  |  |  |  |
| 9 | 6 | 2.10% | 5 | 4.21% | 2009 |  |  |  |  |

#Outliers represented by A/Guizhou/1/2013 and A/Guizhou/2/2013; viruses in this cluster were not reported in 2014.
\*Collection year of last reported sequence or new clade proposal

**Table 2.** New clade descriptions/Candidate vaccine viruses (CVVs)

| Clade | Description of clade | WHO Candidate Vaccine Virus (CVV) |
| --- | --- | --- |
| 2.3.2.1c | China, Japan 2009-2019 | Previously A/duck/Vietnam/NCVD-1584/2012 |
| 2.3.2.1d | China 2014-2016 | A/chicken/Guiyang/1153/2016 |
| 2.3.2.1e | Vietnam, Cambodia, Laos 2012-2024 | A/duck/Vietnam/NCVD-1584/2012, A/Cambodia/SVH240441/2024 |
| 2.3.2.1f | China, India, Middle East, Africa 2013-2018 | A/chicken/Ghana/20/2015 |
| 2.3.2.1g | Indonesia 2012-2023 | no CVV |
| 2.3.4.4 | China 2008-2012 (ancestral viruses to clade 2.3.4.4 will remain 2.3.4.4 with no letter) | no CVV |
| 2.3.4.4a | Vietnam/China A(H5N6) 2013-2015 | A/Sichuan/26221/2014 |
| 2.3.4.4b | 2016/2017 Siberia into Europe/Middle East/Africa/Asia A(H5N8); one human case reported in China, North America 2021-2022 wild birds.<br>North America, Europe, East Asia, 2013-2024 | A/Fujian-Sanyuan/21099/2017; A/Astrakhan/3212/2020; A/Am. Wigeon/South Carolina/22-000345-001/2021; A/Ezo red fox/Hokkaido/1/2022; A/chicken/Ghana/AVL-76321VIR7050-39/2021-like |
| 2.3.4.4c | 2014/2015 Eurasian/North American, intercontinental A(H5N8) and A(H5N2)<br>Korea, Taiwan, United States 2013-2019 | A/gyrfalcon/Washington/41088-6/2014 |
| 2.3.4.4d | Korea/Japan/China A(H5N6); some Chinese A(H5N6) human infections<br>2014-2016 | A/Hubei/29578/2016 |
| 2.3.4.4e | China/Vietnam/Japan/Korea poultry and wild birds and Chinese A(H5N6) human infections<br>2014-2019 | A/duck/Hyogo/1/2016 |
| 2.3.4.4f | China/Vietnam poultry and wild birds<br>2013-2017 | A/chicken/Vietnam/NCVD-15A59/2015 |
| 2.3.4.4g | China/Vietnam 2013-2020 | A/chicken/Vietnam/RAHO4-CD-20-421/2020 |
| 2.3.4.4h | China/Vietnam/Bangladesh/Mongolia 2013-2024 | A/Guangdong/18SF020/2018-like A(H5N6), A/Fujian/2/2024 A(H5N6) |
| 2.3.4.4-like | 2010-2017 outliers | no CVV |

Since the divergence of clade 2.3.4.4 from other fourth-order groups within clade 2.3.4, this lineage has exhibited a high APD of 3.50%, reflecting the rapid emergence, evolution, expansion in host range and geographic expansion of highly divergent A(H5) HA genes (6-12, 17, 23-25). Clade 2.3.4.4 includes viruses with diverse NA subtypes (N1, N2, N3, N5, N6, N7, and N8) and forms multiple distinct monophyletic lineages, which have been subdivided into subclades 2.3.4.4a through 2.3.4.4h. These viruses have been responsible for numerous outbreaks across Asia, the Americas, and Europe in recent years. A(H5N6) viruses detected in China and Vietnam in birds and poultry during 2013-2015 and 2015-2017, respectively, were classified as clade 2.3.4.4a and 2.3.4.4f. Clade 2.3.4.4b, containing N1-N8, spread from East Asia to Africa, the Americas, Antarctica and Europe, and transmitted to multiple mammalian species. Spillover events involving clade 2.3.4.4b viruses, which became the globally dominant clade during this time period, caused multiple detections in wild and domestic mammals, as well as outbreaks in fur-farmed minks and foxes in Europe, dairy cattle in the USA, marine mammals in South America and Antarctica, cats in Europe, North America and Asia, and sporadic human infections following exposure to infected animals. A(H5N2) and A(H5N8) viruses, introduced from East Asia to Europe, Taiwan, Japan and North America through migratory birds, were classified as clade 2.3.4.4c. Clade 2.3.4.4d comprised A(H5N6) viruses detected exclusively in China during 2015–2016. Viruses circulating in East and Southeast Asia from 2013 to 2020 were classified into clades 2.3.4.4e, 2.3.4.4g, and 2.3.4.4h based on phylogeny. Human infections have been reported for the majority of 2.3.4.4 subclades, except 2.3.4.4c and 2.3.4.4f.

Clade 2.3.4.4 (prior to splitting into subclades) had an internal APD of 3.50%, reflecting the rapid evolution of this mixed-subtype group. Furthermore, several of the newly designated 2.3.4.4 subclades had intra-clade APDs over the 1.5% threshold required for further splitting of clades due to the rapid expansion of these clades into new regions (e.g., 1.98% intra-clade APD of 2.3.4.4b representing over 13,000 sequences). Due to sampling biases stemming from the overrepresentation of sequences from specific outbreaks and insufficient data from regions with recent introductions, further subdivision of subclades with intra-clade APDs greater than 1.5% was deferred. Additional genetic data are needed to better assess the endemicity of specific clusters and justify further splits beyond the current subclades 2.3.4.4a–h. Additionally, the division into subclades 2.3.4.4a–h excluded 125 viruses that fell outside these subclades. Many of these outlier viruses, positioned basally within the phylogeny of clade 2.3.4.4, were provisionally designated as “2.3.4.4-like.” The introduction of the “2.3.4.4-like” designation marks a refinement in clade-level classification, reflecting an effort to address the complexity and heterogeneity within rapidly evolving groups like clade 2.3.4.4. This designation was established to account for viruses that do not fit neatly into the defined subclades 2.3.4.4a–h but still share key evolutionary characteristics with clade 2.3.4.4 viruses. Many of these viruses occupy individual positions within the phylogeny and represent early diverging lineages or sequences that exhibit limited clustering with established subclades.

Indeed, continued genetic divergence has been observed in other clades (e.g., 2.1.3.2a), with several exceeding the 1.5% threshold for intra-clade APDs. However, these clades were not further subdivided due to limited circulation in recent years and uncertainty of continued circulation. Finally, no updates were made to clades believed to be “extinct”, as no new sequence data were available for this analysis (Figure 1, Table 1).

## DISCUSSION

Enzootic HPAI A(H5N1) viruses remain a significant global concern, causing recurrent outbreaks in poultry worldwide, and posing risks of zoonotic transmission. Since their initial spread across Eurasia and into North and West Africa, these viruses have become entrenched in poultry populations in geographically isolated areas, causing sporadic outbreaks and human infections (5). Between 2013 and 2019, a resurgence of HPAI activity was observed, largely driven by the emergence of reassortant viruses, including HPAI A(H5N2), A(H5N5), A(H5N6), and A(H5N8), which were detected in Asia, Europe, and later in North America(8, 10). However, since 2021, A(H5N1) has re-emerged as the predominant subtype, primarily due to the widespread global circulation of clade 2.3.4.4b.

The last update from the WHO/WOAH/FAO H5 Evolution Working Group, covering data up to December 31, 2014, identified 13 actively circulating clades during 2013–2014 (5). This nomenclature update proposes the subdivision of clades 2.3.2.1c and 2.3.4.4 into subclades 2.3.2.1c–g and 2.3.4.4a–h, based on their geographic distribution and phylogenetic characteristics. Clade 2.3.2.1c diverged into clade 2.3.2.1d in China and Vietnam in 2014-2016, 2.3.2.1e in Vietnam and Cambodia in 2020-2024, 2.3.2.1f in India, Middle East, and Africa in 2013-2017, and 2.3.2.1g in Indonesia in 2012-2016. These subdivisions highlight the diversification of endemic A(H5) viruses within specific regions and time periods, supported by HA sequence divergence as well as genotype variability among viruses detected globally (data not shown). Meanwhile, clade 2.3.4.4 viruses also exhibit extensive geographic and genetic diversity and have incorporated numerous NA subtypes. The clade’s spread originated in China, extending to Southeast Asia and subsequently diverging into subclades 2.3.4.4a and 2.3.4.4d–h. Transmission events introduced viruses to North America, Taiwan, Europe and Japan forming subclade 2.3.4.4c. Clade 2.3.4.4b expanded from East Asia and Africa to Europe, North America, and South America, leading to outbreaks in multiple mammalian species (16).

To support the adoption of this nomenclature, several bioinformatic tools are publicly available for clade classification of HA sequences with this updated system. These include LABEL (https://github.com/CDCgov/label), Nextstrain (https://nextstrain.org/avian-flu/h5n1/ha/2y) and TIPars (https://tipars.hku.hk/reference/Influenza-A-H5_HA_Tree). These tools are useful and convenient for users to determine which A(H5) clades their new HA sequences belong to, reducing the efforts of bioinformatics processing and to support international uniformity.

In articles published between nomenclature updates, emerging clades such as 2.3.4.4 have been assigned provisional names in the literature that may differ from their formal designation in subsequent revisions. While the WHO/WOAH/FAO H5 Evolution Working Group considers historical convention in its nomenclature revisions, authors are encouraged to add the word *provisional* to describe emerging clades in order to facilitate continuity within the literature in the event that such names are not later adopted by the community at large. Continued global surveillance, monitoring and characterization of HPAI A(H5) viruses from poultry and wild birds, as well as those from mammal and sporadic human infections, will be critical to assess the prevalence and public health significance of these new clades in the future.

## ACKNOWLEDGEMENTS

We thank the H5 Evolution Working Group Members and Collaborators (Supplemental Data S1) for drafting this manuscript and performing sequence and phylogenetic analyses. We acknowledge the laboratories that provided virus samples and sequence data for access to information deposited into the GISAID database; authors and originating/submitting laboratories are listed in Supplementary Data S3. The findings and conclusions in this report are those of the authors and do not necessarily represent the views of the Centers for Disease Control and Prevention or the Agency for Toxic Substances and Disease Registry. This publication contains the collective views of an international group of experts and does not necessarily represent the decisions or the stated policy of the Food and Agriculture Organization of the United Nations (FAO), the World Organisation for Animal Health (WOAH), or the World Health Organization (WHO). ACB and JJ were funded by the UK Department for the Environment, Food and Rural Affairs (Defra) and the devolved Scottish and Welsh governments under grants SV3032, SV3400, SE2227, and SV3006.

## SUPPORTING INFORMATION

**Figure S1 (A, B)**. Phylogenetic relationships of recently diverged A/goose/Guangdong /1/1996 (Gs/GD/96)-like H5 hemagglutinin (HA) genes. Two maximum likelihood trees containing a total of 7,866 HA nucleotide sequences from (Gs/GD/96) lineage HPAI H5 viruses were constructed with 10,000 resamples for local support of topology (above branches) using FastTree2 (GTR+GAMMA). Newly designated clades are annotated and shown in color with the within-clade average pairwise nucleotide distances. Scale bar denotes nucleotide substitutions per site. A (clade 2.3.2.1); B (2.3.4.4)

**Supplementary Data S1**. Members of the World Health Organization/World Organisation for Animal Health/Food and Agriculture Organization (WHO/WOAH/FAO) H5 Evolution Working Group.

### H5 Evolution Working Group Members and Collaborators

The working group was established in 2008 by request of the World Health Organization’s Global Influenza Programme, Department of Epidemic and Pandemic Alert and Response (WHO, GIP, EPR), the World Organization for Animal Health (WOAH, formerly OIE), and the Food and Agriculture Organization (FAO). It currently consists of the following persons:

1. Francesco Bonfante, Istituto Zooprofilattico Sperimentale delle Venezie, Padova, Italy;
2. Ian H. Brown, The Pirbright Institute, Ash Road, Pirbright, Woking, Surrey, UK;
3. Lorcan Carnegie, OFFLU scientist, FAO Rome, Italy
4. Amelia Coggon, OFFLU scientist, FAO Rome, Italy
5. C. Todd Davis, Centers for Disease Control and Prevention, Georgia, USA;
6. Alice Fusaro, Istituto Zooprofilattico Sperimentale delle Venezie, Padova, Italy;
7. Ron A.M. Fouchier, Erasmus University, Netherlands;
8. Yi Guan, The University of Hong Kong, HK SAR, China;
9. Hideki Hasegawa, WHO Collaborating Centre for Reference and Research on Influenza, National Institute of Infectious Diseases, Tokyo, Japan;
10. Yunho Jang, Centers for Disease Control and Prevention, Georgia, USA;
11. Erik A. Karlsson, Virology Unit, Institut Pasteur du Cambodge, Phnom Penh, Cambodia
12. Mia Kim-Torchetti, National Veterinary Services Laboratories, Iowa, USA;
13. Tommy Lam, The University of Hong Kong, HK SAR, China;
14. Nicola S. Lewis, Worldwide Influenza Centre, London, UK;
15. Isabella Monne, Istituto Zooprofilattico Sperimentale delle Venezie, Padova, Italy;
16. Gounalan Pavade, World Organization for Animal Health, Paris, France
17. Magdi Samaan, WHO, GISRS, Geneva, Switzerland
18. Samuel S. Shepard, Centers for Disease Control and Prevention, Atlanta, Georgia, USA;
19. David Suarez, Southeast Poultry Laboratory, USDA, Athens, Georgia, USA;
20. Dayan Wang, Chinese Center for Disease Control and Prevention, Beijing, China;
21. Richard Webby, WHO Collaborating Center for Studies on the Ecology of Influenza in Animals, St. Jude Children’s Research Hospital, Tennessee, USA;
22. Frank Wong, Australian Animal Health Laboratory, Geelong, Australia;
23. Wenqing Zhang, WHO, GISRS, Geneva, Switzerland

**Supplementary Data S2**. List of new sequence data since the previous clade nomenclature update, including viruses, clade designation, gene accession numbers, and database sources for sequences used in the phylogenetic analysis.

**Supplementary Data S3**. List of authors, originating and submitting laboratories of the sequences from GISAID’s EpiFlu™ Database used in this report.

## Notes

* Group members and collaborators are provided in the Supporting Information (Data S1).

### Competing Interest Statement

The authors have declared no competing interest.

