## Supplementary figures and images for "Nomenclature updates to the hemagglutinin gene clade designations resulting from the continued evolution of high pathogenicity avian influenza A(H5) virus clades 2.3.2.1c and 2.3.4.4"

### S. Figure 1A

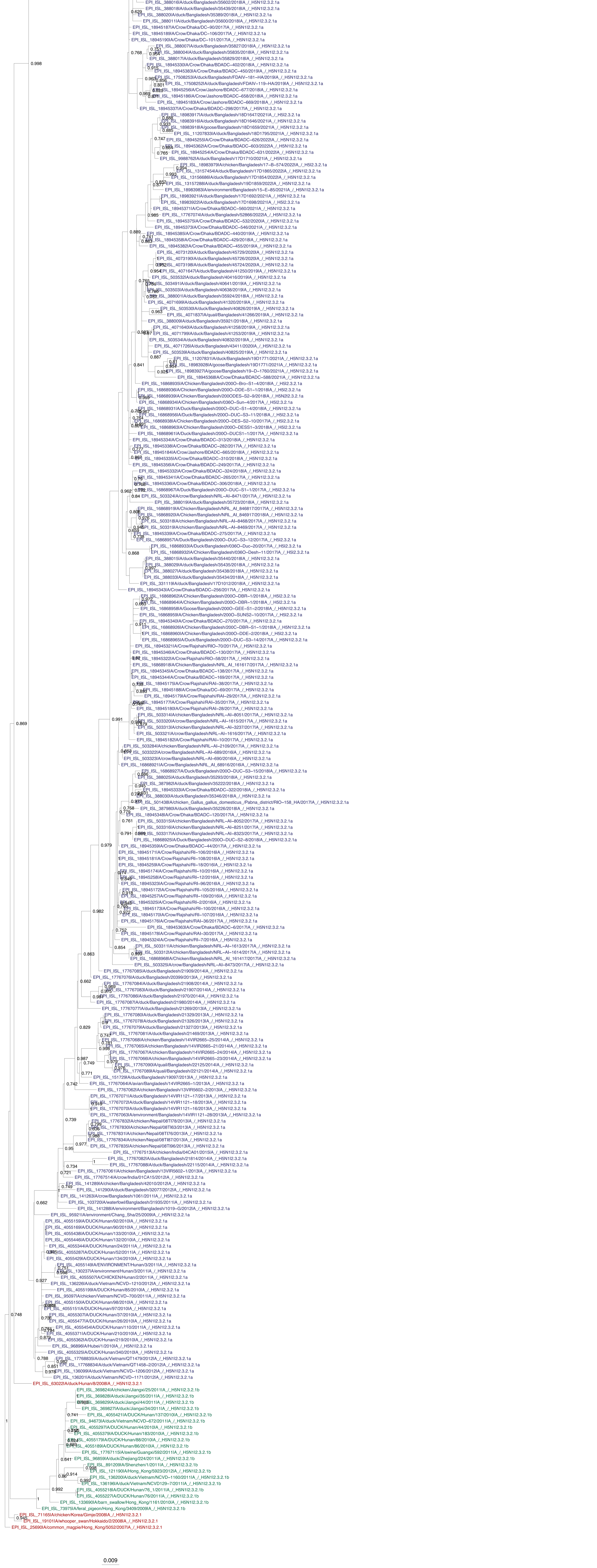

### S. Figure 1B

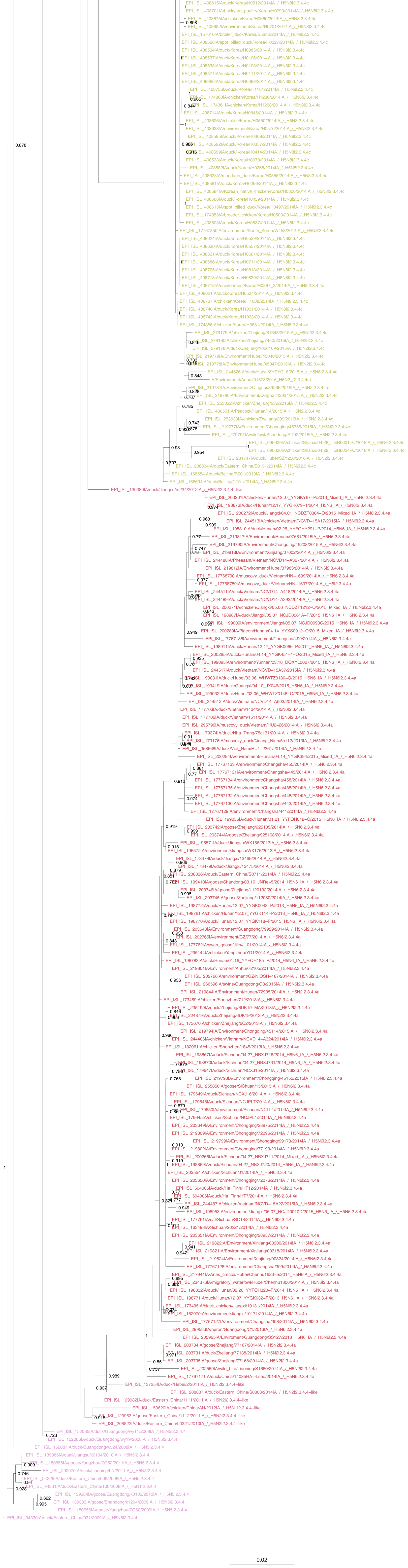
